# DE-STRESS: A user-friendly web application for the evaluation of protein designs

**DOI:** 10.1101/2021.04.28.441790

**Authors:** Michael J. Stam, Christopher W. Wood

## Abstract

*De novo* protein design is a rapidly growing field and there are now many interesting and useful examples of designed proteins in the literature. However, most designs could be classed as failures when characterised in the lab, usually as a result of low expression, misfolding, aggregation or lack of function. This high attrition rate makes protein design unreliable and costly. It is possible that some of these failures could be caught earlier in the design process if it were quick and easy to generate information and a set of high-quality metrics regarding designs, which could be used to make reproducible and data-driven decisions about which designs to characterise experimentally.

We present DE-STRESS (DEsigned STRucture Evaluation ServiceS), a web application for evaluating structural models of designed and engineered proteins. DE-STRESS has been designed to be simple, intuitive to use and responsive. It provides a wealth of information regarding designs, as well as tools to help contextualise the results and formally describe the properties that a design requires to be fit for purpose.

**Availability:** DE-STRESS is available for non-commercial use, without registration, through the following website: https://pragmaticproteindesign.bio.ed.ac.uk/de-stress/. Source code for the application is available on GitHub: https://github.com/wells-wood-research/de-stress. The data used to generate reference sets is available through a GraphQL API, with the following URL: https://pragmaticproteindesign.bio.ed.ac.uk/big-structure/graphql.

## Introduction

There has been rapid development in the field of *de novo* protein design over recent years, with more groups producing increasingly ambitious designs with complex behaviour, often applied in cellular environments (Ben-Sasson *et al*., 2021; Glasgow *et al*., 2019; Harrington *et al*., 2021; Herud-Sikimić *et al*., 2021; Pirro *et al*., 2020; Sesterhenn *et al*., 2020; VanDrisse *et al*., 2021; Pan *et al*., 2020).

Despite the great promise of *de novo* protein design, it remains the domain of highly specialist research groups, as there are significant barriers blocking broader adoption as a methodology. One major challenge is that only a fraction of designs adopt stable, folded structures when expressed (Huang *et al*., 2016), and it can be difficult to identify these models using the metrics calculated during the design process alone (Radom *et al*., 2018). This is especially challenging for designs with complex requirements that are needed for targeted applications.

Here we present DE-STRESS (DEsigned STRucture Evaluation ServiceS), a user-friendly web application for evaluating structural models of designed and engineered proteins. We aim to provide the user with as much information as possible about their designs before they select sequences to characterise experimentally.

## Methods and Results

The DE-STRESS application consists of a simple and intuitive user interface, written in Elm/JavaScript, and a backend web stack, consisting of Gunicorn/Flask/GraphQL/PostgreSQL (figure 1). The interface has three main sections that the user can explore: Designs, Reference Sets and Specifications.

**Figure 1:**
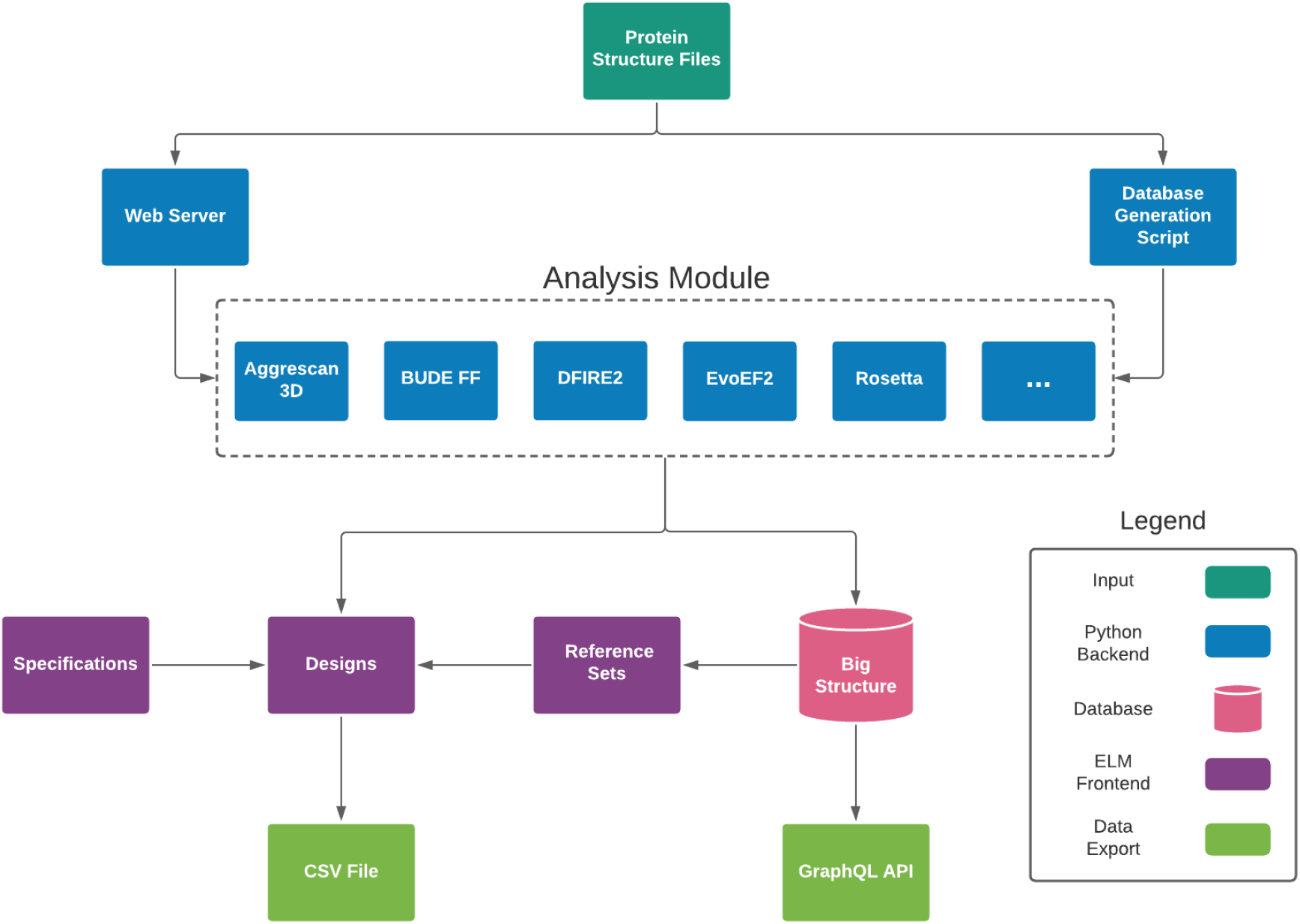
Overview of the DE-STRESS application architecture.

On the Designs page (figure 2A), users can upload models of proteins (in PDB format) to the DE-STRESS server, where all the included metrics will be calculated for each design. Once the metrics have been calculated, an overview of the whole batch of designs is provided. Detailed information can be viewed for each design (figure 2B), as well as a comparison to the active Reference Set and Requirement Specification (*vide infra*).

**Figure 2:**
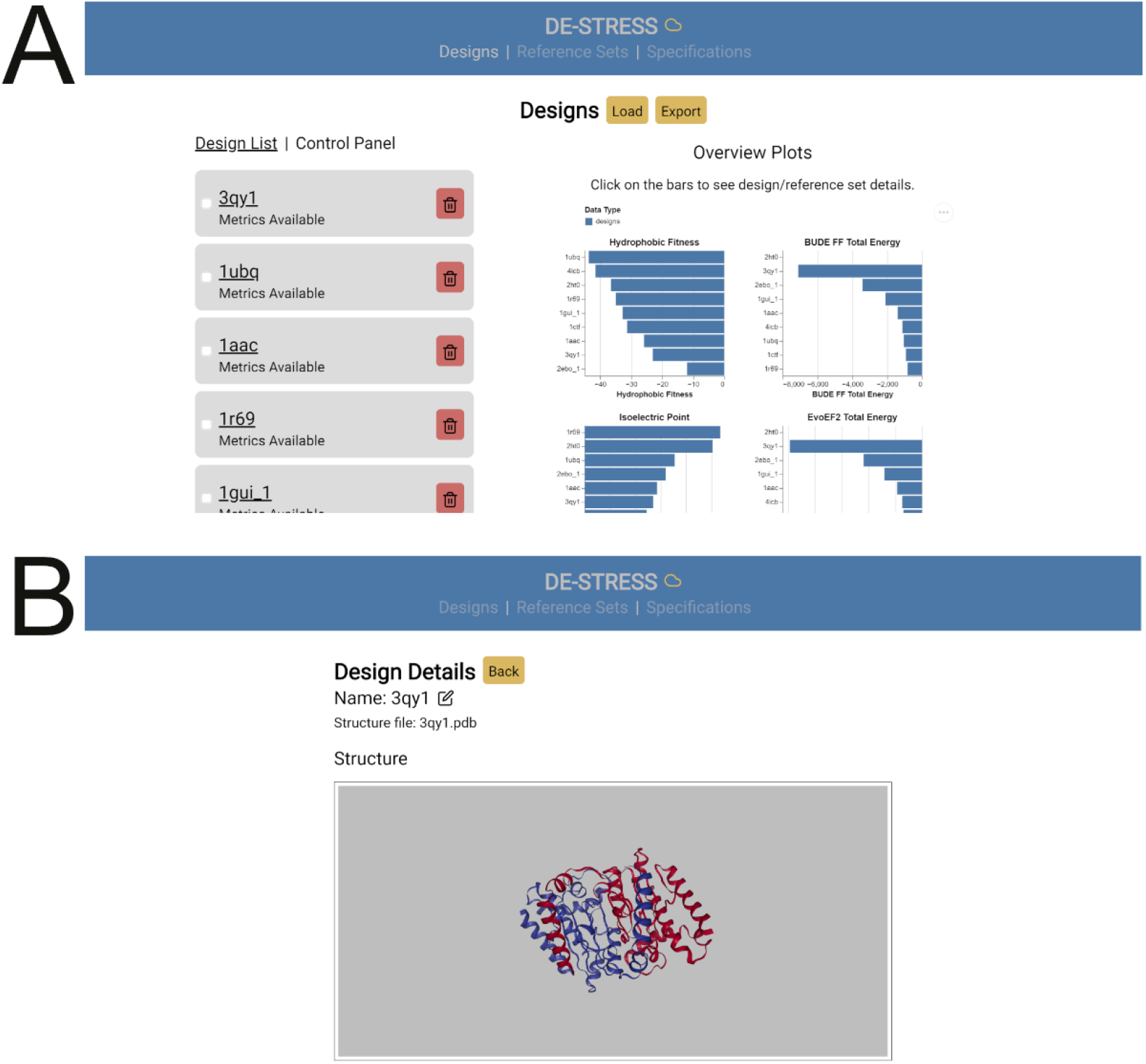
The DE-STRESS user interface. A) The “Designs” page allows users to upload designs, obtain information on the whole batch of designs and download a CSV file containing their results. B) Detailed information on specific designs is offered on the “Design Details” page.

On the Reference Sets page, users can define a set of known protein structures from the PDB (Berman *et al*., 2003), which can be used as a basis of comparison for their designs. We have precalculated the metrics included in DE-STRESS for the biological units, as defined by the PDBe (http://ftp.ebi.ac.uk/pub/databases/pdb/data/biounit/), of 82,010 protein structures. The remaining structures in the PDB either did not contain protein, contained formatting errors in the PDB file or, in the case of large structures, failed to return results within a reasonable timeframe. Similar restrictions have been placed on models that can be uploaded to the webserver, in order to ensure server stability. Using these data, the user can define their own reference sets by submitting a list of PDB accession codes, enabling them to compare their designs to relevant structures. Additionally, two default reference sets are provided as an example, based on high-quality structures from Top500 (Hobohm and Sander, 1994) and Pisces (Wang and Dunbrack, 2003). Finally, users can also define reference sets from models that they have uploaded to the server, allowing the creation of reference sets from unpublished structures.

Once a reference set has been defined, aggregated metrics are presented alongside the metrics for the user’s designs. All the data used to generate the reference sets are available to search and download, programmatically and interactively, through a GraphQL API available at the following url: https://pragmaticproteindesign.bio.ed.ac.uk/big-structure/graphql.

Reference sets are intended to provide context for the metrics returned by DE-STRESS, and while we have provided some example reference sets of high-quality crystal structures, manually defining a reference set is much more useful. For example, if you are designing coiled-coil proteins, comparing your designs to similar structures is likely to be more useful than a general set of proteins, as the metrics and other information, such as sequence composition, will be directly comparable. Furthermore, it is worth noting that designs with metrics that fall outside of the range observed for known protein structures may still express, fold and be functional. In certain instances, the designer may be actively selecting designs with metrics that fall outside of those observed for natural proteins, in fact this is very often the purpose of designing proteins *de novo*.

The Specifications page allows the user to define “Requirement Specifications”, which encapsulate the properties their designs should have in order to be fit for purpose. The user can define complex rules, such as nested conditional properties, that can be used to filter designs, alongside associated metadata. We plan to expand the role of the specifications in the future, allowing the user to capture more information about their design intent and export the specification to be used by other programmes.

A variety of external software packages are used by the DE-STRESS web server to calculate metrics for uploaded models (table 1, supplementary table 1), with detailed and up-to-date information on the version and command used to run the software provided in the supplementary material and glossary page of the application. Basic information is extracted using ISAMBARD (Wood *et al*., 2017), such as the isoelectric point and composition of the sequence, as well as implementations of metrics from the literature, such as packing density (Weiss, 2007) and hydrophobic fitness (Huang *et al*., 1995). In addition to these metrics, DE-STRESS applies a range of scoring functions that have either been developed specifically for protein design or have been applied to design proteins, such as BUDE FF (McIntosh-Smith *et al*., 2012, 2015; Thomson *et al*., 2014), EvoEF2 (Huang *et al*., 2020b, 2, 2020a), Rosetta (Alford *et al*., 2017; Cao *et al*., 2020) and DFIRE2 (Yang and Zhou, 2008; Negron and Keating, 2014). Finally, we use Aggrescan3D (Kuriata *et al*., 2019) to calculate an aggregation propensity score for protein structures, as aggregation is a common failure mode for designed and engineered proteins (Marques *et al*., 2021). For detailed information regarding the metrics included see supplementary tables 2-11. These metrics are presented on the Design Details page, alongside a visualisation of the model, using the NGL JavaScript library (Rose and Hildebrand, 2015; Rose *et al*., 2016), and other information such as secondary structure assignment using DSSP (Kabsch and Sander, 1983; Touw *et al*., 2015).

**Table 1:** Model evaluation methods included in DE-STRESS. The scoring convention indicates whether the score is considered more favourable if it is lower (-ve) or higher (+ve).

| Evaluation Method | Type | Scoring Convention | Reference |
| --- | --- | --- | --- |
| **Packing Density** | Geometric Analysis | +ve | (Weiss, 2007) |
| **Hydrophobic Fitness** |  | -ve | (Huang et al., 1995) |
| **BUDE FF** | All-Atom Scoring Function | -ve | (McIntosh-Smith et al., 2012, 2015) |
| **EvoEF2** |  | -ve | (Huang et al., 2020, 2) |
| **Rosetta** |  | -ve | (Alford et al., 2017) |
| **DFIRE2** | Statistical Potential | -ve | (Yang and Zhou, 2008) |
| **Aggrescan3D** | Aggregation Propensity | -ve | (Kuriata et al., 2019) |
| **ISAMBARD** | Basic Information | N/A | (Wood et al., 2017) |
| **DSSP** | Secondary-Structure Assignment | N/A | (Kabsch and Sander, 1983; Touw et al., 2015) |

A privacy first approach has been taken when implementing DE-STRESS. No login is required to use the application and no data regarding the user, or their designs, are stored on our server. Designs are submitted directly to an in-memory job queue, with no associated metadata, and the results are returned directly to the user. All data regarding the user’s designs are stored locally on the device used to access the website and can be exported to a CSV file for further analysis. With this architecture, we aim to give the user confidence in submitting their designs to the server. However, if they would like to take further steps to ensure that no one could access their data, they can run a local instance of the web application, which we have made as simple as possible by containerising the application.

We envisage that, for many users, DE-STRESS will be useful for generating descriptive information and statistics that could be manually examined to choose designs that meet the needs of their application. Beyond this, the datasets that DE-STRESS creates could be useful for automatic identification of high-quality models using data-driven methods. As a simple example of this, we attempted to discern experimentally-determined structures from decoys that are used to benchmark protein-folding algorithms. The dataset and the associated scripts for performing this analysis are available on GitHub: https://github.com/wells-wood-research/stam-m-wood-c-de-stress-2021.

Using the DE-STRESS web application, we generated and exported metrics for a random sample of 9 experimentally-determined structures, along with 360 decoys (40 per structure) from a set that was previously generated by 3DRobot. 3DRobot produces structurally diverse and well-packed decoy structures that are difficult to discern from experimentally-determined structures with simple metrics, such as percentage of secondary structure or radius of gyration, as well as more complex metrics such as statistical potentials (Deng et al., 2016). We also included alternative structures for each of these proteins (supplementary table 14), identified through the structural similarity search tool on the RSCB PDB, to ensure that the findings were robust to minor variations in the structure.

Firstly, before performing PCA, various metrics were excluded from the data set. Categorical and discrete variables were excluded as PCA requires continuous variables, and metrics that were constant for a structure were excluded. This is because these metrics provided no information that could be used to distinguish between the decoys and experimentally-determined structures. In addition to this, as the energy values are dependent on the size of the protein structure, we normalised for length by dividing all the energy values by the number of residues in the structure. A full list of the metrics included in the analysis is shown in the supplementary table 12. Before performing PCA, the metrics were normalised using min-max normalisation. The first two principal components, which explained 60% of the variance in the data, were plotted against each other to explore how well the DE-STRESS metrics differentiated the experimentally-determined structures from the decoys (figure 3). The additional crystallographic structures were also included in these plots.

**Figure 3:**
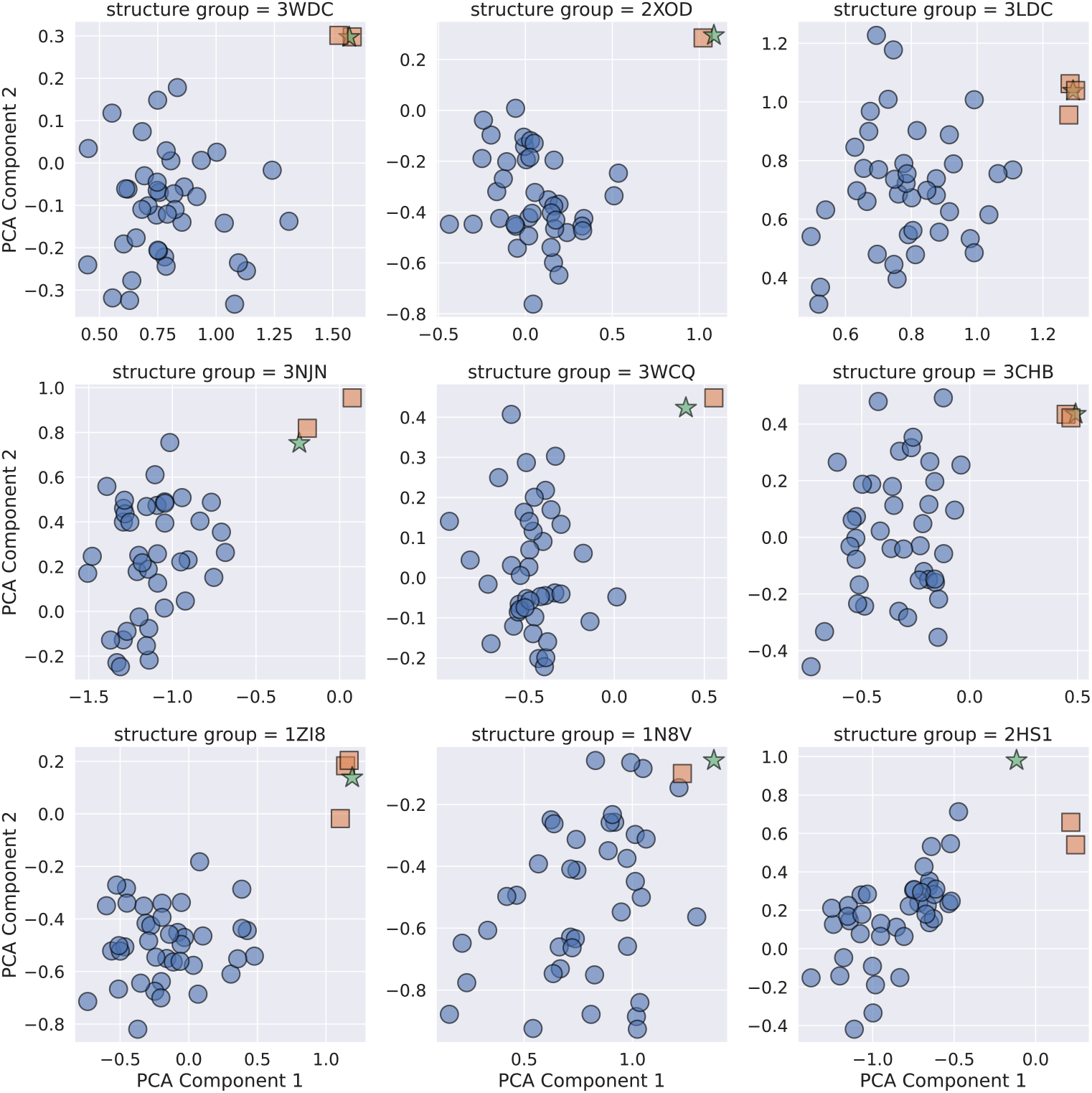
Principal component analysis of DE-STRESS metrics generated for experimentally-determined structures (stars), folding decoys (circles) and alternative crystallographic structures of the protein (squares).

From the plots in figure 3, we see that the experimentally-determined structures, shown as stars, are generally distinct from the decoys and usually have the largest values for both principal components 1 and 2. In addition to this, the additional crystallographic structures, shown as squares, are close to the experimentally-determined structures used to generate the 3DRobot set. This analysis suggests that the DE-STRESS metrics are capturing properties of the structural models that can be used to distinguish “native-like” structures from decoys.

To understand which metrics differentiated the experimentally-determined structures from the decoys, we examined the relative contribution of individual DE-STRESS metrics to the first two principal components (supplementary table 13). It was found that the major contributors to PC1 were the short range and long-range hydrogen bonding terms in from the Rosetta forcefield and the total BUDE forcefield energy value. The major contributors to PC2 were the average score of Aggrescan3D, the steric component of the BUDE forcefield and a solvation term from the Rosetta forcefield. Long-range hydrogen bonding was highlighted by the developers of 3DRobot as a key differentiating feature between native structures and decoys, and maintaining this property was a key to creating native-like decoys. However, while analysis performed in the 3DRobot publications shows that their decoys performed significantly better in this regard than previous decoy sets (Deng *et al*., 2016), our results indicate that there is still some detectable difference between the experimentally-determined structures and the decoys. The fact that Aggrescan3D is a key contributor to PC2 might indicate that the surface properties of the decoys are significantly different from the native structures, and this could indicate a potential route to improve the process used to create decoy models.

## Conclusions

DE-STRESS enables both non-experts and seasoned protein designers to rapidly evaluate their designs, providing a framework for making reproducible, data-driven decisions about which design to take forward for experimental characterisation. While some protocols and applications have been developed to meet similar needs as DE-STRESS (Bernhofer et al., 2021; Guffy et al., 2018; Yallapragada et al., 2020, 2), none of them have the same breadth of metrics and tools, all packaged in a user-friendly web application. However, the metrics included in the initial version of DE-STRESS are certainly not exhaustive: general methods for assessing the quality of a protein model (Vriend, 1990; Wang *et al*., 2019; Williams *et al*., 2018; Laskowski *et al*., 1993), as well as more specialised methods for assessing design quality, such as analysis of covariation similarity with homologs (Ollikainen and Kortemme, 2013; Ó Conchúir *et al*., 2015; Ludwiczak *et al*., 2018), could be useful additions in future releases. We believe that our analysis of decoy structures demonstrates that the metrics included in this initial version of DE-STRESS are useful and can be used to identify high-quality models, but further experimental work is required to determine whether this translates into a reduced failure rate of designs taken into the lab, which would meet our ultimate aim of increasing the efficiency of the protein-design process and making it more accessible and reliable as a technique.

## Acknowledgements

The authors would like to thank Lynne Regan, Dek Woolfson for feedback on the manuscript/application and Kathryn Shelly for feedback on the application. We would also like to thank the UoE School of Biological Sciences IT department for infrastructure support, and the developers of the software used to generate many of the metrics included in DE-STRESS.

