## Supplementary Data for "DE-STRESS: A user-friendly web application for the evaluation of protein designs"

| Programme name | Description | Command Used | Citations |
|---|---|---|---|
| Aggrescan3D 2.0 (v1.0.2) | Aggregation Propensity | aggrescan -i pdb_file_path -w output -D 10 -v 4 | Kuriata et al. (2019). Aggrescan3D standalone package for structure-based prediction of protein aggregation properties. Bioinformatics 35, 3834–3835. |
| BUDE (v1.0.0) | Energy Function | import ampal import budeff design = ampal.load_pdb( pdb_string, path=False) budeff.get_internal_energy( design)) | McIntosh-Smith et al. (2012). Benchmarking Energy Efficiency, Power Costs and Carbon Emissions on Heterogeneous Systems. The Computer Journal 55, 192–205. McIntosh-Smith et al. (2015). High performance in silico virtual drug screening on many-core processors. The International Journal of High Performance Computing Applications 29, 119–134. |

| | | | |
|---|---|---|---|
| DFIRE2-pair | Energy Function | calene dfire_pair.lib pdb_file_path | Yang et al. (2008). Ab initio folding of terminal segments with secondary structures reveals the fine difference between two closely related all-atom statistical energy functions. Protein Science 17, 1212–1219. |
| DSSP (v2.0.4) | Secondary Structure Assignment | import isambard.evaluation as ev import ampal design=ampal.load_pdb(pdb_string, path=False) ev.tag_dssp_data(design) sequence_info={ chain.id: SequenceInfo( sequence="".join(m.mol_letter for m in chain), dssp_assignment="".join( m.tags["dssp_data"]["ss_definition"] for m in chain ), ) for chain in design if isinstance(chain, ampal.Polypeptide) } | Kabsch et al. (1983). Dictionary of protein secondary structure: Pattern recognition of hydrogen-bonded and geometrical features. Biopolymers 22, 2577–2637. Touw et al. (2015). A series of PDB-related databanks for everyday needs. Nucleic Acids Research 43, D364–D368. |
| EvoEF2 (EvoEF v2) | Energy Function | EvoEF2 --command = ComputeStability --pdb = pdb_file_path | Huang et al. (2020). EvoEF2: accurate and fast energy function for computational protein design. Bioinformatics 36, 1135–1142. |
| Hydrophobic Fitness (ISAMBARD v2.3.1) | Energy Function | import isambard.evaluation as ev import ampal design=ampal.load_pdb(pdb_string, path=False) ev.calculate_hydrophobic_fitness( design) | Huang et al. (1995). Recognizing native folds by the arrangement of hydrophobic and polar residues. J Mol Biol 252, 709–720. Wood et al. (2017). ISAMBARD: an open-source computational environment for biomolecular analysis, modelling and design. Bioinformatics 33, 3043–3050. |

| | | | |
|---|---|---|---|
| Packing Density(ISAMBARD v2.3.1) | Geometric Analysis | import isambard.evaluation as ev import ampal design=ampal.load_pdb(pdb_string, path=False) ev.tag_packing_density(design) mean_packing_density = np.mean( [a.tags["packing density"] for a in design.get_atoms() if a.element != "H"]) | Weiss (2007). On the interrelationship between atomic displacement parameters (ADPs) and coordinates in protein structures. Acta Crystallogr D Biol Crystallogr 63, 1235–1242. Wood et al. (2017). ISAMBARD: an open-source computational environment for biomolecular analysis, modelling and design. Bioinformatics 33, 3043–3050. |
| Rosetta (ref2015) | Energy Function | rosetta_src_2020.08.61146 _bundle/main/source/bin/score _jd2.linuxgccrelease -in:file:s pdb_file_path - ignore_unrecognized_res - scorefile_format json | Alford et al. (2017). The Rosetta All-Atom Energy Function for Macromolecular Modeling and Design. J. Chem. Theory Comput. 13, 3031–3048. |

*Table S1: This table provides a description of the programmes that are included in DE-STRESS, the command used by DE-STRESS to obtain the results and the citations.*

| **Metric Name** | **Metric Description** | **Variable Name in CSV Output** |
|---|---|---|
| Total Score | This value is a global indicator of the aggregation propensity/solubility of the protein structure. It depends on the protein size. It allows assessing changes in solubility promoted by amino acid substitutions in a particular protein structure. The more negative the value, the highest the global solubility. | aggrescan3d: total_value |
| Average Score | This value is a normalized indicator of the aggregation propensity/solubility of the protein structure. Allows comparing the solubility of different protein structures. It also allows assessing changes in solubility promoted by amino acid substitutions in a particular protein structure. The more negative the value, the highest the normalized solubility. | aggrescan3d: avg_value |
| Minimum Score | This is the value of the most soluble residue in the structural context. | aggrescan3d: min_value |
| Maximum Score | This is the value of the most aggregation-prone residue in the structural context. | aggrescan3d: max_value |

*Table S2: This table provides a description of the metrics from the Aggrescan3D 2.0 programme and their variable names in the CSV output file.*

| Metric Name | Metric Description | Variable Name in CSV Output |
|---|---|---|
| Total Energy | This value is the total BUDE force field energy. It is the sum of the steric, desolvation and charge components. | budeff: total |
| Steric Energy | This value is the steric component of the BUDE force field energy. It is calculated with a simplified Leonard-Jones potential. It is softer than the steric component of many other force fields. | budeff: steric |
| Desolvation Energy | This value is the desolvation component of the BUDE force field energy. | budeff: desolvation |
| Charge Energy | This value is the charge component of the BUDE force field energy. | budeff: charge |

*Table S3: This table provides a description of the metrics from the BUDE programme and their variable names in the CSV output file.*

| Metric Name | Metric Description | Variable Name in CSV Output |
|---|---|---|
| Total Energy | This value is the total DFIRE2 energy. This is the only field that is returned from running DFIRE2 on a pdb file. | dfire2 - total |

*Table S4: This table provides a description of the metrics from the DFIRE2 programme and their variable names in the CSV output file.*

| Metric Name | Metric Description | Variable Name in CSV Output |
|---|---|---|
| H | α-helix | Not included in csv output |
| B | Isolated β-bridge | Not included in csv output |
| E | Extended β-strand | Not included in csv output |
| G | 3-10 helix | Not included in csv output |
| I | π-helix | Not included in csv output |
| T | Hydrogen-bonded turn | Not included in csv output |
| S | Bend | Not included in csv output |
| - | Loop | Not included in csv output |

*Table S5: This table provides a description of the metrics from the DSSP programme and their variable names in the CSV output file.*

| Metric Name | Metric Description | Variable Name in CSV Output |
|---|---|---|
| Total Energy | This value is the total EvoEF2 energy. It is the sum of the reference, intra residue, inter residue - same chain and inter residue - different chains, energy values. In the EvoEF2 output this field is called `Total`. | evoef2: total |
| Reference Energy | This value is the total reference energy. This value is not included in the EvoEF2 output and is calculated in DE-STRESS. | evoef2: ref total |
| Intra Residue Energy | This value is the total energy for intra residue interactions. This value is not included in the EvoEF2 output and is calculated in DE-STRESS. | evoef2: intraR total |

| | | |
|---|---|---|
| Inter Residue - Same Chain Energy | This value is the total energy for inter residue interactions in the same chain. This value is not included in the EvoEF2 output and is calculated in DE-STRESS. | evoef2: interS total |
| Inter Residue - Different Chains Energy | This value is the total energy for inter residue interactions in different chains. This value is not included in the EvoEF2 output and is calculated in DE-STRESS. | evoef2 - interD total |
| ALA - Reference | This value is reference energy for the amino acid Alanine (ALA). In the EvoEF2 output this value is called \`reference_ALA\`. | Not included in csv output |
| ARG - Reference | This value is reference energy for the amino acid Arginine (ARG). In the EvoEF2 output this value is called \`reference_ARG\`. | Not included in csv output |
| ASN - Reference | This value is reference energy for the amino acid Asparagine (ASN). In the EvoEF2 output this value is called \`reference_ASN\`. | Not included in csv output |
| ASP - Reference | This value is reference energy for the amino acid Aspartic acid (ASP). In the EvoEF2 output this value is called \`reference_ASP\`. | Not included in csv output |
| CYS - Reference | This value is reference energy for the amino acid Cysteine (CYS). In the EvoEF2 output this value is called \`reference_CYS\`. | Not included in csv output |
| GLN - Reference | This value is reference energy for the amino acid Glutamine (GLN). In the EvoEF2 output this value is called \`reference_GLN\`. | Not included in csv output |
| GLU - Reference | This value is reference energy for the amino acid Glutamic acid (GLU). In the EvoEF2 output this value is called \`reference_GLU\`. | Not included in csv output |
| GLY - Reference | This value is reference energy for the amino acid glycine (GLY). In the EvoEF2 output this value is called \`reference_GLY\`. | Not included in csv output |
| HIS - Reference | This value is reference energy for the amino acid Histidine (HIS). In the EvoEF2 output this value is called \`reference_HIS\`. | Not included in csv output |
| ILE - Reference | This value is reference energy for the amino acid Isoleucine (ILE). In the EvoEF2 output this value is called \`reference_ILE\`. | Not included in csv output |
| LEU - Reference | This value is reference energy for the amino acid Leucine (LEU). In the EvoEF2 output this value is called \`reference_LEU\`. | Not included in csv output |
| LYS - Reference | This value is reference energy for the amino acid Lysine (LYS). In the EvoEF2 output this value is called \`reference_LYS\`. | Not included in csv output |
| MET - Reference | This value is reference energy for the amino acid Methionine (MET). In the EvoEF2 output this value is called \`reference_MET\`. | Not included in csv output |
| PHE - Reference | This value is reference energy for the amino acid Phenylalanine (PHE). In the EvoEF2 output this value is called \`reference_PHE\`. | Not included in csv output |
| PRO - Reference | This value is reference energy for the amino acid Proline (PRO). In the EvoEF2 output this value is called \`reference_PRO\`. | Not included in csv output |
| SER - Reference | This value is reference energy for the amino acid Serine (SER). In the EvoEF2 output this value is called \`reference_SER\`. | Not included in csv output |
| THR - Reference | This value is reference energy for the amino acid Threonine (THR). In the EvoEF2 output this value is called \`reference_THR\`. | Not included in csv output |

| | | |
|---|---|---|
| TRP - Reference | This value is reference energy for the amino acid Tryptophan (TRP). In the EvoEF2 output this value is called `reference_TRP`. | Not included in csv output |
| TYR - Reference | This value is reference energy for the amino acid Tyrosine (TYR). In the EvoEF2 output this value is called `reference_TYR`. | Not included in csv output |
| VAL - Reference | This value is reference energy for the amino acid Valine (VAL). In the EvoEF2 output this value is called `reference_VAL`. | Not included in csv output |
| VDW Attractive - Intra Residue | This value is the Van der Waals attractive energy for intra residue interactions. In the EvoEF2 output this value is called `intraR_vdwatt`. | Not included in csv output |
| VDW Repulsive - Intra Residue | This value is the Van der Waals repulsive energy for intra residue interactions. In the EvoEF2 output this value is called `intraR_vdwrep`. | Not included in csv output |
| Electrostatics - Intra Residue | This value is the Coulomb's electrostatics energy for intra residue interactions. In the EvoEF2 output this value is called `intraR_electr`. | Not included in csv output |
| Desolvation Polar - Intra Residue | This value is the polar atoms desolvation energy for intra residue interactions. In the EvoEF2 output this value is called `intraR_deslvP`. | Not included in csv output |
| Desolvation Non Polar - Intra Residue | This value is the non polar atoms desolvation energy for intra residue interactions. In the EvoEF2 output this value is called `intraR_deslvH`. | Not included in csv output |
| HB Sidechain Backbone Distance - Intra Residue | This value is the energy for the hydrogen-acceptor distance from sidechain - backbone and intra residue interactions. In the EvoEF2 output this value is called `intraR_hbscbb_dis`. | Not included in csv output |
| HB Sidechain Backbone Theta - Intra Residue | This value is the energy for the angle between the donor, hydrogen and acceptor atoms (theta), from sidechain - backbone and intra residue interactions. In the EvoEF2 output this value is called `intraR_hbscbb_the`. | Not included in csv output |
| HB Sidechain Backbone Phi - Intra Residue | This value is the energy for the angle between the hydrogen, acceptor and base atoms (phi), from sidechain - backbone and intra residue interactions. In the EvoEF2 output this value is called `intraR_hbscbb_phi`. | Not included in csv output |
| Amino Acid Propensity - Intra Residue | This value is the amino acid propensity energy for intra residue interactions. In the EvoEF2 output this value is called `aapropensity`. | Not included in csv output |
| Ramachandran - Intra Residue | This value is the Ramachandran energy for intra residue interactions. In the EvoEF2 output this value is called `ramachandran`. | Not included in csv output |
| Dunbrack Rotamer - Intra Residue | This value is the Dunbrack Rotamer energy for intra residue interactions. In the EvoEF2 output this value is called `dunbrack`. | Not included in csv output |
| VDW Attractive - Inter Residue - Same Chain | This value is the Van der Waals attractive energy for inter residue interactions - same chain. In the EvoEF2 output this value is called `interS_vdwatt`. | Not included in csv output |
| VDW Repulsive - Inter Residue - Same Chain | This value is the Van der Waals repulsive energy for inter residue interactions - same chain. In the EvoEF2 output this value is called `interS_vdwrep`. | Not included in csv output |
| Electrostatics - Inter Residue - Same Chain | This value is the Coulomb's electrostatics energy for inter residue interactions - same chain. In the EvoEF2 output this value is called `interS_electr`. | Not included in csv output |
| Desolvation Polar - Inter Residue - Same Chain | This value is the polar atoms desolvation energy for inter residue interactions - same chain. In the EvoEF2 output this value is called `interS_deslvP`. | Not included in csv output |

| | | |
|---|---|---|
| Desolvation Non Polar - Inter Residue - Same Chain | This value is the non polar atoms desolvation energy for inter residue interactions - same chain. In the EvoEF2 output this value is called `interS_deslvH`. | Not included in csv output |
| Disulfide Bonding - Inter Residue - Same Chain | This value is the disulfide bonding energy for inter residue interactions - same chain. In the EvoEF2 output this value is called `interS_ssbond`. | Not included in csv output |
| HB Backbone Backbone Distance - Inter Residue - Same Chain | This value is the energy for the hydrogen-acceptor distance from backbone - backbone and inter residue interactions - same chain. In the EvoEF2 output this value is called `interS_hbbbbb_dis`. | Not included in csv output |
| HB Backbone Backbone Theta - Inter Residue - Same Chain | This value is the energy for the angle between the donor, hydrogen and acceptor atoms (theta), from backbone - backbone and inter residue interactions - same chain. In the EvoEF2 output this value is called `interS_hbbbbb_the`. | Not included in csv output |
| HB Backbone Backbone Phi - Inter Residue - Same Chain | This value is the energy for the angle between the hydrogen, acceptor and base atoms (phi), from backbone - backbone and inter residue interactions - same chain. In the EvoEF2 output this value is called `interS_hbbbbb_phi`. | Not included in csv output |
| HB Sidechain Backbone Distance - Inter Residue - Same Chain | This value is the energy for the hydrogen-acceptor distance from side chain - backbone and inter residue interactions - same chain. In the EvoEF2 output this value is called `interS_hbscbb_dis`. | Not included in csv output |
| HB Sidechain Backbone Theta - Inter Residue - Same Chain | This value is the energy for the angle between the donor, hydrogen and acceptor atoms (theta), from side chain - backbone and inter residue interactions - same chain. In the EvoEF2 output this value is called `interS_hbscbb_the`. | Not included in csv output |
| HB Sidechain Backbone Phi - Inter Residue - Same Chain | This value is the energy for the angle between the hydrogen, acceptor and base atoms (phi), from side chain - backbone and inter residue interactions - same chain. In the EvoEF2 output this value is called `interS_hbscbb_phi`. | Not included in csv output |
| HB Sidechain Sidechain Distance - Inter Residue - Same Chain | This value is the energy for the hydrogen-acceptor distance from side chain - side chain and inter residue interactions - same chain. In the EvoEF2 output this value is called `interS_hbscsc_dis`. | Not included in csv output |
| HB Sidechain Sidechain Theta - Inter Residue - Same Chain | This value is the energy for the angle between the donor, hydrogen and acceptor atoms (theta), from side chain - side chain and inter residue interactions - same chain. In the EvoEF2 output this value is called `interS_hbscsc_the`. | Not included in csv output |
| HB Sidechain Sidechain Phi - Inter Residue - Same Chain | This value is the energy for the angle between the hydrogen, acceptor and base atoms (phi), from side chain - side chain and inter residue interactions - same chain. In the EvoEF2 output this value is called `interS_hbscsc_phi`. | Not included in csv output |
| VDW Attractive - Inter Residue - Different Chains | This value is the Van der Waals attractive energy for inter residue interactions - different chains. In the EvoEF2 output this value is called `interD_vdwatt`. | Not included in csv output |
| VDW Repulsive - Inter Residue - Different Chains | This value is the Van der Waals repulsive energy for inter residue interactions - different chains. In the EvoEF2 output this value is called `interD_vdwrep`. | Not included in csv output |
| Electrostatics - Inter Residue - Different Chains | This value is the Coulomb’s electrostatics energy for inter residue interactions - different chains. In the EvoEF2 output this value is called `interD_electr`. | Not included in csv output |

| | | |
|---|---|---|
| Desolvation Polar - Inter Residue - Different Chains | This value is the polar atoms desolvation energy for inter residue interactions - different chains. In the EvoEF2 output this value is called `interD_deslvP`. | Not included in csv output |
| Desolvation Non Polar - Inter Residue - Different Chains | This value is the non polar atoms desolvation energy for inter residue interactions - different chains. In the EvoEF2 output this value is called `interD_deslvH`. | Not included in csv output |
| Disulfide Bonding - Inter Residue - Different Chains | This value is the disulfide bonding energy for inter residue interactions - different chains. In the EvoEF2 output this value is called `interD_ssbond`. | Not included in csv output |
| HB Backbone Backbone Distance - Inter Residue - Different Chains | This value is the energy for the hydrogen-acceptor distance from backbone - backbone and inter residue interactions - different chains. In the EvoEF2 output this value is called `interD_hbbbbb_dis`. | Not included in csv output |
| HB Backbone Backbone Theta - Inter Residue - Different Chains | This value is the energy for the angle between the donor, hydrogen and acceptor atoms (theta), from backbone - backbone and inter residue interactions - different chains. In the EvoEF2 output this value is called `interD_hbbbbb_the`. | Not included in csv output |
| HB Backbone Backbone Phi - Inter Residue - Different Chains | This value is the energy for the angle between the hydrogen, acceptor and base atoms (phi), from backbone - backbone and inter residue interactions - different chains. In the EvoEF2 output this value is called `interD_hbbbbb_phi`. | Not included in csv output |
| HB Sidechain Backbone Distance - Inter Residue - Different Chains | This value is the energy for the hydrogen-acceptor distance from side chain - backbone and inter residue interactions - different chains. In the EvoEF2 output this value is called `interD_hbscbb_dis`. | Not included in csv output |
| HB Sidechain Backbone Theta - Inter Residue - Different Chains | This value is the energy for the angle between the donor, hydrogen and acceptor atoms (theta), from side chain - backbone and inter residue interactions - different chains. In the EvoEF2 output this value is called `interD_hbscbb_the`. | Not included in csv output |
| HB Sidechain Backbone Phi - Inter Residue - Different Chains | This value is the energy for the angle between the hydrogen, acceptor and base atoms (phi), from side chain - backbone and inter residue interactions - different chains. In the EvoEF2 output this value is called `interD_hbscbb_phi`. | Not included in csv output |
| HB Sidechain Sidechain Distance - Inter Residue - Different Chains | This value is the energy for the hydrogen-acceptor distance from side chain - side chain and inter residue interactions - different chains. In the EvoEF2 output this value is called `interD_hbscsc_dis`. | Not included in csv output |
| HB Sidechain Sidechain Theta - Inter Residue - Different Chains | This value is the energy for the angle between the donor, hydrogen and acceptor atoms (theta), from side chain - side chain and inter residue interactions - different chains. In the EvoEF2 output this value is called `interD_hbscsc_the`. | Not included in csv output |
| HB Sidechain Sidechain Phi - Inter Residue - Different Chains | This value is the energy for the angle between the hydrogen, acceptor and base atoms (phi), from side chain - side chain and inter residue interactions - different chains. In the EvoEF2 output this value is called `interD_hbscsc_phi`. | Not included in csv output |

*Table S6: This table provides a description of the metrics from the EvoEF2 programme and their variable names in the CSV output file.*

| **Metric Name** | **Metric Description** | **Variable Name in CSV Output** |
|---|---|---|
| Hydrophobic Fitness | This value is an efficient centroid-based method for calculating the packing quality of the protein structure. For | hydrophobic fitness |

| | |
|---|---|
| | this method C, F, I, L, M, V, W and Y are considered hydrophobic. |

*Table S7: This table provides a description of the metrics from the Hydrophobic Fitness programme and their variable names in the CSV output file.*

| **Metric Name** | **Metric Description** | **Variable Name in CSV Output** |
|---|---|---|
| Isoelectric Point | This value is the pH of a solution at which the net charge of the protein becomes zero. | isoelectric point (pH) |

*Table S8: This table provides a description of the metrics from the Isoelectric Point programme and their variable names in the CSV output file.*

| **Metric Name** | **Metric Description** | **Variable Name in CSV Output** |
|---|---|---|
| Packing Density | This value is an efficient centroid-based method for calculating the packing quality of the protein structure. For this method C, F, I, L, M, V, W and Y are considered hydrophobic. | packing density |

*Table S9: This table provides a description of the metrics from the Packing Density programme and their variable names in the CSV output file.*

| **Metric Name** | **Metric Description** | **Variable Name in CSV Output** |
|---|---|---|
| Total Energy | This value is the total Rosetta energy. It is a weighted sum of the different Rosetta energy values. In the Rosetta `score.sc` output file, this value is called `total_score`. | rosetta - total |
| Reference | This value is the reference energy for the different amino acids. In the Rosetta `score.sc` output file, this value is called `ref`. | Not included in csv output |
| VDW Attractive | This value is the attractive energy between two atoms on different residues separated by distance, d. In the Rosetta `score.sc` output file, this value is called `fa_atr`. | rosetta - fa_atr |
| VDW Repulsive | This value is the repulsive energy between two atoms on different residues separated by distance, d. In the Rosetta `score.sc` output file, this value is called `fa_rep`. | rosetta - fa_rep |
| VDW Repulsive Intra Residue | This value is the repulsive energy between two atoms on the same residue separated by distance, d. In the Rosetta `score.sc` output file, this value is called `fa_intra_rep`. | rosetta - fa_intra_rep |
| Electrostatics | This value is the energy of interaction between two non-bonded charged atoms separated by distance, d. In the Rosetta `score.sc` output file, this value is called `fa_elec`. | rosetta - fa_elec |
| Solvation Isotropic | This value is the Gaussian exclusion implicit solvation energy between protein atoms in different residues. In the Rosetta `score.sc` output file, this value is called `fa_sol`. | rosetta - fa_sol |
| Solvation Anisotropic Polar Atoms | This value is the orientation-dependent solvation of polar atoms assuming ideal water geometry. In the Rosetta `score.sc` output file, this value is called `lk_ball_wtd`. | rosetta - lk_ball_wtd |

| | | |
|---|---|---|
| Solvation Isotropic Intra Residue | This value is the Gaussian exclusion implicit solvation energy between protein atoms in the same residue. In the Rosetta `score.sc` output file, this value is called `fa_sol_intraR`. | rosetta - fa_intra_sol_xover4 |
| HB Long Range Backbone | This value is the energy of long range hydrogen bonds. In the Rosetta `score.sc` output file, this value is called `hbond_lr_bb`. | rosetta - hbond_lr_bb |
| HB Short Range Backbone | This value is the energy of short range hydrogen bonds. In the Rosetta `score.sc` output file, this value is called `hbond_sr_bb`. | rosetta - hbond_sr_bb |
| HB Backbone Sidechain | This value is the energy of backbone-side chain hydrogen bonds. In the Rosetta `score.sc` output file, this value is called `hbond_bb_sc`. | rosetta - hbond_bb_sc |
| HB Sidechain Sidechain | This value is the energy of side chain-side chain hydrogen bonds. In the Rosetta `score.sc` output file, this value is called `hbond_sc`. | rosetta - hbond_sc |
| Disulfide Bridges | This value is the energy of disulfide bridges. In the Rosetta `score.sc` output file, this value is called `dslf_fa13`. | rosetta - dslf_fa13 |
| Backbone Torsion Preference | This value is the probability of backbone $\phi$, $\psi$ angles given the amino acid type. In the Rosetta `score.sc` output file, this value is called `rama_prepro`. | rosetta - rama_prepro |
| Amino Acid Propensity | This value is the probability of amino acid identity given the backbone $\phi$, $\psi$ angles. In the Rosetta `score.sc` output file, this value is called `p_aa_pp`. | rosetta - p_aa_pp |
| Dunbrack Rotamer | This value is the probability that a chosen rotamer is native-like given backbone $\phi$, $\psi$ angles. In the Rosetta `score.sc` output file, this value is called `fa_dun`. | rosetta - fa_dun |
| Omega Penalty | This value is a backbone-dependent penalty for cis $\omega$ dihedrals that deviate from 0° and trans $\omega$ dihedrals that deviate from 180°. In the Rosetta `score.sc` output file, this value is called `omega`. | rosetta - omega |
| Open Proline Penalty | This value is a penalty for an open proline ring and proline $\omega$ bonding energy. In the Rosetta `score.sc` output file, this value is called `pro_close`. | rosetta - pro_close |
| Tyrosine $\chi$3 Dihedral Angle Penalty | This value is a sinusoidal penalty for non-planar tyrosine $\chi$3 dihedral angle. In the Rosetta `score.sc` output file, this value is called `yhh_planarity`. | rosetta - yhh_planarity |

*Table S10: This table provides a description of the metrics from the Rosetta programme and their variable names in the CSV output file.*

| **Metric Name** | **Metric Description** | **Variable Name in CSV Output** |
|---|---|---|
| ALA - Composition | The proportion of residues that are Alanine (ALA) in the structure. | composition: ALA |
| ARG - Composition | The proportion of residues that are Arginine (ARG) in the structure. | composition: ARG |
| ASN - Composition | The proportion of residues that are Asparagine (ASN) in the structure. | composition: ASN |
| ASP - Composition | The proportion of residues that are Aspartic Acid (ASP) in the structure. | composition: ASP |
| CYS - Composition | The proportion of residues that are Cysteine (CYS) in the structure. | composition: CYS |
| GLN - Composition | The proportion of residues that are Glutamine (GLN) in the structure. | composition: GLN |
| GLU - Composition | The proportion of residues that are Glutamic Acid (GLU) in the structure. | composition: GLU |

| | | |
|---|---|---|
| GLY - Composition | The proportion of residues that are Glycine (GLY) in the structure. | composition: GLY |
| HIS - Composition | The proportion of residues that are Histidine (HIS) in the structure. | composition: HIS |
| ILE - Composition | The proportion of residues that are Isoleucine (ILE) in the structure. | composition: ILE |
| LEU - Composition | The proportion of residues that are Leucine (LEU) in the structure. | composition: LEU |
| LYS - Composition | The proportion of residues that are Lysine (LYS) in the structure. | composition: LYS |
| MET - Composition | The proportion of residues that are Methionine (MET) in the structure. | composition: MET |
| PHE - Composition | The proportion of residues that are Phenylalanine (PHE) in the structure. | composition: PHE |
| PRO - Composition | The proportion of residues that are Proline (PRO) in the structure. | composition: PRO |
| SER - Composition | The proportion of residues that are Serine (SER) in the structure. | composition: SER |
| THR - Composition | The proportion of residues that are Threonine (THR) in the structure. | composition: THR |
| TRP - Composition | The proportion of residues that are Tryptophan (TRP) in the structure. | composition: TRP |
| TYR - Composition | The proportion of residues that are Tyrosine (TYR) in the structure. | composition: TYR |
| VAL - Composition | The proportion of residues that are Valine (VAL) in the structure. | composition: VAL |
| UNK - Composition | The proportion of residues that are Unknown (UNK) in the structure. | composition: UNK |
| Number of Residues | The number of amino acid residues in the structure. | number of redidues |
| Mass (Da) | The mass of the structure in daltons (Da). | mass (da) |

*Table S11: This table provides a description of other metrics included in DE-STRESS and their variable names in the CSV output file.*

| **Included Metrics** | **Excluded Metrics** |
|---|---|
| aggrescan3d: avg_value | aggrescan3d: total_value |
| aggrescan3d: min_value | composition: ALA |
| aggrescan3d: max_value | composition: ARG |
| budeff: total | composition: ASN |
| budeff: steric | composition: ASP |
| budeff: desolvation | composition: CYS |
| budeff: charge | composition: GLN |
| dfire2 - total | composition: GLU |
| evoef2: total | composition: GLY |
| evoef2: intraR total | composition: HIS |
| evoef2: interS total | composition: ILE |
| hydrophobic fitness | composition: LEU |
| packing density | composition: LYS |
| rosetta - total | composition: MET |
| rosetta - fa_atr | composition: PHE |
| rosetta - fa_rep | composition: PRO |

| | |
|---|---|
| rosetta - fa_intra_rep | composition: SER |
| rosetta - fa_elec | composition: THR |
| rosetta - fa_sol | composition: TRP |
| rosetta - lk_ball_wtd | composition: TYR |
| rosetta - fa_intra_sol_xover4 | composition: VAL |
| rosetta - hbond_lr_bb | composition: UNK |
| rosetta - hbond_sr_bb | design name |
| rosetta - hbond_bb_sc | decoy or native |
| rosetta - hbond_sc | evoef2 - interD total |
| rosetta - rama_prepro | evoef2: ref total |
| rosetta - p_aa_pp | isoelectric point (pH) |
| rosetta - fa_dun | mass (da) |
| rosetta - omega | number of residues |
| rosetta - pro_close | pdb id |
| | rosetta - dslf_fa13 |
| | rosetta - yhh_planarity |
| | structure group |

*Table S12: This table lists the metrics that were included in the decoy analysis and those that were excluded.*

| **Top 6 contributors to PC1** | **Top 6 contributors to PC2** |
|---|---|
| rosetta - hbond_sr_bb | aggrescan3d: avg_value |
| rosetta - hbond_lr_bb | budeff: steric |
| budeff: total | rosetta - fa_intra_sol_xover4 |
| evoef2: intraR total | aggrescan3d: max_value |
| budeff: charge | aggrescan3d: min_value |
| dfire2 - total | packing density |

*Table S13: This table shows the top 6 DE-STRESS metrics that contribute to Principal Component 1 (PC1) and Principal Component 2 (PC2).*

| **Experimentally-determined structure** | **Additional crystallographic structures** |
|---|---|
| 1N8V A | 1N8U A |
| 1ZI8 A | 1ZIC A, 1ZIX A, 1ZI9 A |
| 2HS1 A | 2HS2 A, 3S53 A |
| 2XOD A | 2X2P A |
| 3CHB D | 1PZK D, 1PZJ D |
| 3LDC A | 3OUS A, 3R65 A, 3LDD A |
| 3NJN A | 3NJH A, 3NJM A |
| 3WCQ A | 3AB5 A |
| 3WDC A | 3WDE A, 3WDD A |

*Table S14: This table shows the pdb ids and chains for the additional structures that were selected from the PDB, and the experimentally determined structures from the 3DRobot_set.*
